# Estrogen-related receptor alpha promotes skeletal muscle regeneration and mitigates muscular dystrophy

**DOI:** 10.1101/2025.05.15.654390

**Authors:** Thi Thu Hao Nguyen, Ya Xiang Huang, Svitlana Poliakova, Citu Citu, Eira Mann, Danesh H. Sopariwala, Zhongming Zhao, Ashok Kumar, Vihang A. Narkar

## Abstract

Skeletal muscle regeneration in chronic muscle diseases such as Duchenne Muscular Dystrophy (DMD) has remained clinically unsurmountable. Estrogen-related receptor alpha (ERRα) plays a critical role in adult skeletal muscle metabolism and exercise fitness. Whether ERRα activation can drive muscle regeneration and mitigation of dystrophy in DMD is not known. We have investigated ERRα signaling in pre-clinical models of acute muscle injury and DMD. ERRα is induced in differentiating C2C12 myoblast and regenerating muscle. ERRα silencing suppressed proliferation and differentiation in C2C12 myoblasts. RNA sequencing revealed that angiogenic factor and proliferation genes were downregulated by ERRα knockdown in proliferating cells, whereas oxidative mitochondrial and differentiation regulator genes were downregulated in differentiating cells. In accordance with in vitro findings, transgenic ERRα overexpression in rodent skeletal muscle stimulates muscle regeneration after acute BaCl_2_ injury, which is accompanied by enhanced angiogenesis and mitochondrial biogenesis. Notably, ERRα and its angiogenic and metabolic target gene expression is suppressed in muscle stem cells (MuSCs) derived from dystrophic muscles in mdx mice, coinciding with proliferation and differentiation defect in these cells. Loss of ERRα and its target gene expression was recapitulated in adult dystrophic mdx muscles. Consequently, muscle specific ERRα overexpression in mdx mice restored angiogenic and metabolic gene expression, induced vascular and oxidative remodeling, alleviated baseline muscle damage, and boosted regeneration after BaCl2 injury in dystrophic muscle. Our studies demonstrate a pro-regenerative role of ERRα and its deficiency in dystrophic muscles and its MuSCs. ERRα restoration could be a therapeutic strategy for DMD through angio-metabolic gene program.

## 1. INTRODUCTION

Skeletal muscle homeostasis is maintained through its plasticity to adapt in myofiber type, size and regenerative capacity in response to environmental cues such as inactivity, exercise and nutrition, as well as in aging and myopathy. The regeneration of skeletal muscle is attributed primarily to muscle stem cells (MuSCs). In response to muscle injury, MuSCs are activated and undergo several rounds of proliferation and then differentiate into myoblasts, which fuse with each other or injured myofibers to accomplish muscle repair [1; 2]. Muscle diseases (e.g. muscular dystrophies) and other chronic conditions (e.g. diabetes) decrease muscle regenerative capacity, subsequently impairing muscle repair and homeostasis [3–6]. Specifically, in Duchenne Muscular Dystrophy (DMD) repeated cycles of degeneration-regeneration result in MuSC exhaustion and scar tissue formation [7]. While key growth factors (e.g. Igf and Fgf) and regulators (e.g. Pax7 and MyoD) have been elucidated in skeletal muscle regeneration, so far they haven’t been translated to effective muscle therapy [8–10]. Unraveling new mechanisms of muscle regeneration may have future impact on regenerative therapies for muscle wasting diseases.

Estrogen-related receptors (ERRα, β, γ) are nuclear receptors and transcriptional regulators of mitochondrial biogenesis, oxidative metabolism, gluconeogenesis and thermogenesis in adulthood [11–13]. ERRα and ERRγ are highly expressed in the skeletal muscle. ERRβ expression is undetectable at protein level in the skeletal muscle. Using muscle specific knockout mice, we and others recently showed that muscle ERRα and ERRγ are individually dispensable and compensatory; but together are indispensable for mitochondrial biogenesis, muscle capillarization, and exercise tolerance in mice [14; 15]. Additional support for ERRs in muscle homeostasis comes from our studies using muscle specific ERRα (TG) transgenic mice, where overexpression of ERRα in muscle predominantly drives an oxidative metabolic and paracrine angiogenesis gene program [16; 17]. Indeed, skeletal muscles of ERRα TG mice show increased oxidative mitochondrial capacity, type IIA myofibers, and capillary density [16; 17]. Furthermore, muscle-specific ERRα overexpression promotes robust neo-angiogenesis and vascular regeneration in response to hindlimb ischemia via paracrine muscle-to-endothelial cell signaling [16; 17]. By contrast, muscle-specific ERRα knockout study demonstrates that exercise-induced muscle angiogenic factors and capillarization require intact muscle ERRα expression [18]. While ERRα is ubiquitously expressed in all muscle types, ERRγ is predominantly expressed in oxidative muscle [19]. Nevertheless, it is increasingly evident that ERRγ has similar angiogenic and metabolic effects in the adult skeletal muscle as ERRα [19–22].

Despite clear link between ERRs and adult muscle aerobic fitness, ERRs remain poorly explored in myopathies. Notably, ERRα expression is repressed in various models of muscular dystrophy [23–25]. ERRα expression is also repressed in clinical muscular dystrophy biopsies [26–28]. While these and other published studies [29; 30] point to a role of ERRα in muscle regeneration, effect of activating ERRα on muscle regeneration in chronic degenerative myopathies such as DMD is unknown. Furthermore, transcriptional mechanisms through which ERRα regulates muscle precursor cell function and myofiber regeneration are poorly understood. In this study, we report that ERRα stimulates proliferation and differentiation of myogenic cells and skeletal muscle regeneration in response to acute injury and in dystrophic muscle of mdx mice (a model of DMD) via angiogenic and metabolic (referred to as angio-metabolic) gene programming.

## 2. METHODS

### 2.1. Mice

Skeletal muscle-specific ERRα overexpressing transgenic mice (TG), in which ERRα expression is under the control of the human skeletal alpha-actin (HSA) promoter, were previously generated in our laboratory [31]. Age-matched TG and WT male mice were used for muscle injury and regeneration at 8-12 weeks of age. Mdx and C57Bl/6J mice were obtained from Jackson Labs. The homozygous female mdx mice were crossed with ERRα TG mice to obtain muscle specific ERRα mdx transgenic mice (mdx-TG) and littermate mdx mice. All mice were maintained, and studies performed as per the U.S. National Institute of Health Guide for Care and Use of Laboratory Animals guidelines, and the procedures were approved by the Animal Welfare Committee at The University of Texas Health Science Center in Houston.

### 2.2. BaCl_2_ skeletal muscle injury model

Muscle injury was induced in WT, TG, mdx, and mdx-TG mice. To induce muscle injury, 20 µl of BaCl_2_ (1.2% in saline) was injected into the tibialis anterior (TA) muscle of each hindlimb in the mice. The day of injury was considered as 0 days post-injury (dpi). Mice were euthanized at 4, 7, and 15 dpi, and TA muscles were collected for further analysis.

### 2.3. Serum creatine kinase (CK) activity

CK was measured in blood collected from the tail vein. Blood was allowed to clot for one hour and centrifuged at 13,000 rpm for 20 min. Serum was collected, and CK activity was measured using a Pointe Liquid Creatine Kinase reagent set.

### 2.4. Cell culture

Mouse C2C12 myoblasts (CRL1772 from ATCC, Manassas, VA) were cultured in growth medium containing DMEM with 10% fetal bovine serum (FBS) and 1% penicillin-streptomycin until 70-80% confluence. Cells were detached from culture plates with 0.25% trypsin and sub-cultured for experiments. Muscle stem cells were isolated from hindlimb muscles by Magnetic-Activated Cell Sorting (MACS) from mdx and non-mdx mice using satellite cell isolation kit and Anti-Integrin α-7 MicroBead (Miltenyi Biotec). For differentiation, myoblasts or muscle stem cells were seeded into a 24-well plate at a density of 80% confluence, and the medium was changed to differentiation medium containing DMEM with 2% heat-inactivated horse serum and 1% penicillin-streptomycin after 24 hours.

### 2.5. Cell proliferation assay

C2C12 or muscle stem cells were seeded at low density in a 96-well plate and cultured in a growth medium. Cell proliferation rate was measured using The Cell Proliferation Kit I (MTT) (Roche, Seoul, Korea). Briefly, 10 µl of MTT reagent was added to each well with 100 µl growth medium, and the cells were incubated for 4 hours at 37°C in a 5% CO_2_ incubator. Then, 100 µl of solubilization buffer was added to each well, and the plate was incubated overnight at 37°C in a 5% CO_2_ incubator. The absorbance was measured at 570 nm with 670 nm as reference wavelength using a microplate reader.

### 2.6. Histology

TA muscles were dissected, frozen in liquid nitrogen-cooled isopentane, and cryo-sectioned at 10 µm thickness. Histological analysis was performed using hematoxylin and eosin staining followed by sequential dehydration with 70%, 95%, and 100% ethanol and clearing in Histoclear solution. Then, the slides were mounted using a Vectamount permanent mounting medium.

### 2.7. Immunostaining

For fluorescent antibody staining, cryosections were fixed and permeabilized with ice-cold methanol before blocking with 10% goat serum in Phosphate Buffer Saline (PBS) for 1 hour at room temperature. Cryosections were incubated overnight with primary antibodies at 4°C. Myoblasts were fixed with 3.7% formaldehyde, permeabilized with 1% NP-40, blocked with 10% goat serum, and incubated with primary antibodies overnight at 4°C. Primary antibodies used included eMyHC (F1.652, Developmental Studies Hybridoma Bank-DSHB, 3 µg/ml), MyoG (F5D, DSHB, 3 µg/ml), MF-20 (DSHB, 3 µg/ml), Laminin (L9393, Sigma, 1:100), CD31 (MCA2388, Bio-Rad, 1:100), and SDHB (ab14714, Abcam, 3 µg/ml). Sections and cells were then washed with PBS before incubation at room temperature for 2 hours with Cy3 or Cy5, Alexa Fluor 488 conjugated goat/donkey anti-mouse or anti-rabbit and anti-rat IgG secondary antibodies (Jackson Immuno Research Laboratories,1:400). For EdU assay, cells were immunolabeled using Click-iT^®^ EdU Imaging Kit (Invitrogen) according to manufacturer’s protocol. Nuclei were counterstained with DAPI or Hoechst, and images were acquired using fluorescence microscopy (Nikon).

### 2.8. Immunoblotting

TA, and C2C12, or muscle stem cells were homogenized in RIPA buffer (Thermo Fisher Scientific) and centrifuged at 13000 g for 15 min at 4°C. Total protein concentration was determined using Pierce BCA protein assay kit (Thermo Fisher Scientific). Proteins were resolved on SDS-PAGE and transferred to PVDF membranes. The membrane was blocked with 3% BSA, 0.1% Tween-20, and Tri’s buffer saline (TBS) for 1 hour at room temperature. The primary antibody diluted with 3% BSA, 0.1% Tween-20, and TBS was incubated overnight at 4°C with continuous shaking. The primary antibodies and dilution factors used include ERRα (13826, Cell signaling, 1:1000), ERRγ (kindly provided by Ron Evans, 1:500), eMyHC (F1.652, DSHB, 0.3 µg/ml), MyoG (F5D, DSHB, 0.3 µg/ml), MF-20 (DSHB, 0.3 µg/ml), Pax7 (DSHB, 0.3 µg/ml), MyoD (sc760, Santa Cruz Biotechnology, 1:500), Vinculin (13901, Cell signaling, 1:1000), NGFR (p75NTR, 8238, Cell signaling, 1:1000), HGF (MA5-14160, Thermo Fisher Scientific, 1:500), VEGFA (sc507, Santa Cruz Biotechnology, 1:500), FGF1 (MA5-35690, Thermo Fisher Scientific, 1:1000), total OXPHOS rodent antibody cocktail (ab110413, Abcam, 1:1000), TFAM (8076, Cell signaling, 1:1000), CYCS (sc13561, Santa Cruz Biotechnology, 1:500). The membranes were washed two times for 40 min with TBST (0.1% Tween-20, TBS) and incubated with secondary horseradish peroxidase-conjugated antibody (Cell signaling, 1:5000) for 2 hours at room temperature. Subsequently, proteins were visualized by chemiluminescence.

### 2.9. siRNA knockdown

Cells were seeded in triplicate at the desired density and incubated for 6 hours before being transfected with *Esrra,* or control scrambled siRNAs. Solutions containing smart pool *Esrra* siRNA or control scrambled siRNA and RNAiMAX were incubated at room temperature for 10 min and added into each well in growth medium for 24 hours at 37°C and 5% CO_2_.

### 2.10. RNA extraction and qRT-PCR

Total RNA was extracted from cells or homogenized TA with TRIzol reagent and Purelink Kit (Ambion, Life Technologies, Carlsbad, CA). RNA was transcribed into cDNA using SuperScript™ III Reverse Transcriptase (Thermo Fisher Scientific). Selected genes were amplified and detected using the Applied Biosystems^TM^ PowerUp^TM^ SYBR^TM^ Green Master Mix (Thermo Fisher Scientific) with a Bio-Rad XF96 cycler (Bio-Rad). Relative gene expression was calculated by normalizing to *Ppia* (*Cyclophilin)* or *Actb* reference genes. All qPCR mouse primers are listed in **Supplemental Table 1**.

### 2.11. RNA Sequencing and differential gene expression analysis

RNA sequencing (RNA-seq) was conducted on total RNA extracted from cultured C2C12 proliferating myoblasts and differentiated myotubes. RNA-seq library was prepared using enriched poly (A)-tailed mRNA in the UT Cancer Genomics Center. The trimming of FASTQ reads to remove the low-quality reads and adapter sequences were done using fastp (v 0.23.2), and quality assessment of the trimmed FASTQ reads was conducted using FastQC (https://www.bioinformatics.babraham.ac.uk/projects/fastqc) [32]. Subsequently, the trimmed reads were aligned on mouse genome reference (GRCm39) using STAR (v2.7.10b) [33]. After alignment, *featureCounts* from the Subread package (v2.0.6) was used to generate gene count matrix, which represented the number of reads corresponds to each gene [34]. Differential expression analysis was performed with DESeq2 to identify upregulated and downregulated genes between knockdown and control groups in C2C12 proliferating myoblasts and differentiated myotube cell lines, with absolute log2FC >0.58 and a q-value cut-off < 0.05 [35]. Here, a log2FC > 0.58 corresponds to 1.5-fold change in gene expression, representing the magnitude of differential expression between conditions. To dissect the profile of each group, the list of differentially expressed genes (DEGs) and entire detectable genes derived from each group were used for GO enrichment analysis and pathway enrichment analysis. We used clusterProfiler version 4.6.2 and followed the standard procedure with the RNA-seq dataset. The enriched GO terms (BP, MF, and CC) were identified using the enrichGO function in this package using parameters (pAdjustMethod = ”BH, pvalueCutoff = 0.05, qvalueCutoff = 0.05). The enriched pathways from KEGG were identified using the enrichKEGG function in clusterProfiler using parameters (pAdjustMethod = ”BH, pvalueCutoff = 0.05, qvalueCutoff = 0.05). The top 30 GO terms (if found) or top 30 pathways (if found) were plotted using the dotplot function in enrichplot_1.18.4 R package [36].

### 2.12. Statistical analysis

Statistical analysis was performed with GraphPad Prism. In all graphs, data are presented as the mean ± standard error of the mean (SEM). Data was analyzed using One-way ANOVA followed by Turkey’s post-hoc test to determine the significance of multiple groups and unpaired Student’s *t*-test between two groups. P-values lower than 0.05 are considered statistically significant.

## 3. RESULTS

### 3.1. ERRα is required for proliferation and differentiation of cultured myoblasts

We first investigated the role of ERRα on the proliferation and differentiation of cultured myoblasts, and myocellular transcriptomic regulation. C2C12 myoblasts were transfected with ERRα (gene name: *Esrra*) siRNA (si*Esrra*) followed by measurement of cellular proliferation. There was a significant reduction in the proliferation of *Esrra* KD myoblasts compared to controls transfected with scrambled siRNA (siNT) **(Figure 1A)**. We also examined EdU incorporation rate for cells in the S phase of the cell cycle. Consistent with cell growth kinetics, EdU incorporation into the replicating DNA was significantly decreased in *Esrra* KD myoblasts compared to the controls **(Figure 1B, C)**.

**Figure 1.**
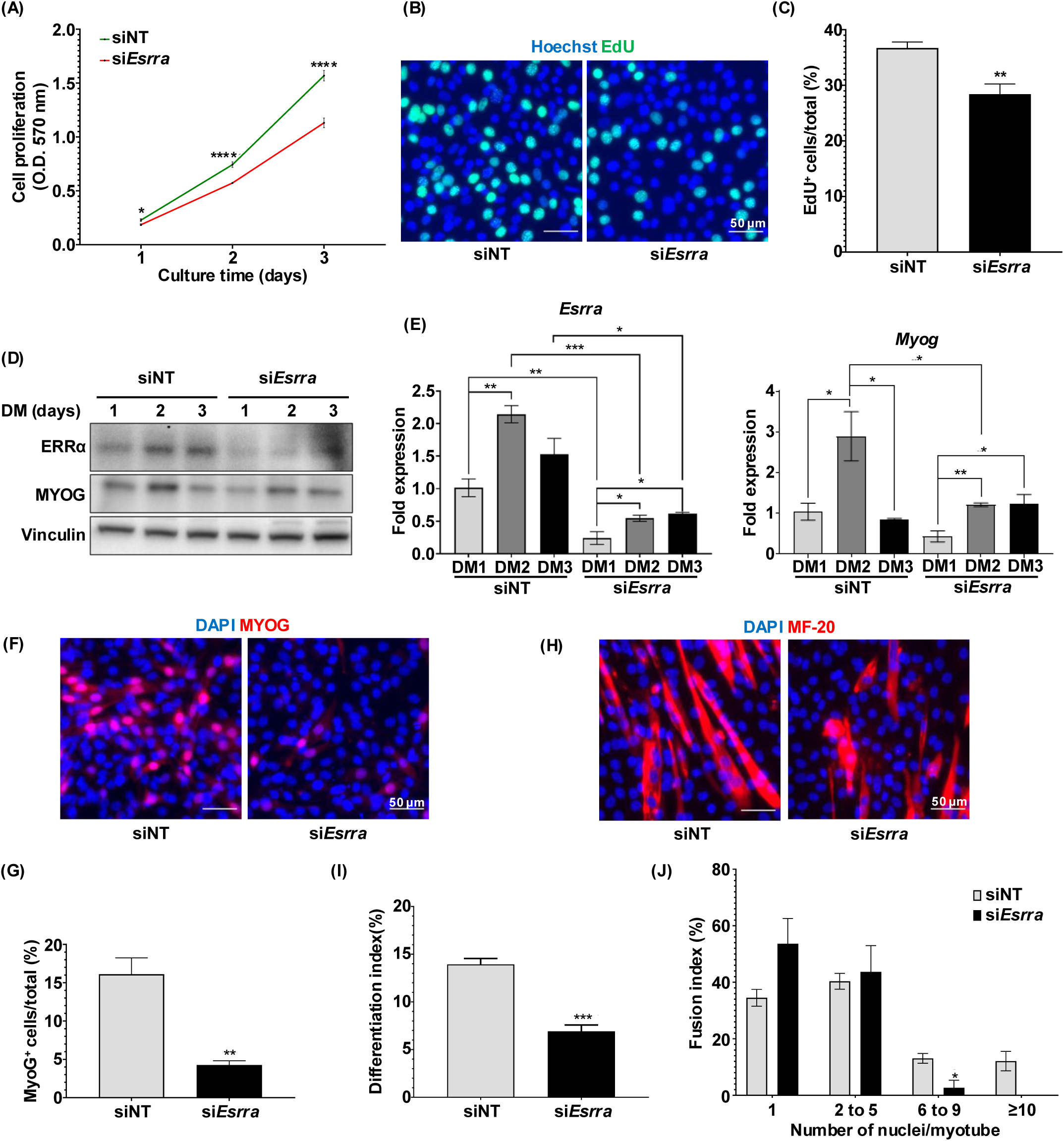
ERRα is required for myoblast proliferation and myogenic differentiation. Following measurements were made in C2C12 cells treated with control scramble siRNA (siNT) or *Esrra* siRNA (si*Esrra*). **(A)** Proliferation rate in C2C12 myoblasts (*p<0.05, ****p<0.0001, N=8). **(B)** Immunofluorescence staining of S-phase cells (identified as EdU^+^cells) in proliferating C2C12 cells (N=5). Scale bar = 50 µm. **(C)** The percentage of S-phase cells (EdU^+^ cells) to total cell number (**p<0.01, N=5). **(D)** Representative western blot (WB) of ERRα and Myogenin (MyoG) expression during myoblast differentiation (N=3). **(E)** mRNA expression of *Esrra* and *Myog* in differentiating C2C12 cells (*p<0.05, **p<0.01, ***p<0.001, N=3). **(F)** Immunofluorescence staining of early differentiated cells (MyoG^+^ cells) at day 1 after myogenic differentiation (N=4). Scale bar = 50 µm. **(G)** Quantification of MyoG^+^ cells to total cell ratio (**p<0.01, N=4). **(H)** Immunofluorescence staining of myotubes (MF20^+^ myotube) at day 3 after myogenic differentiation (N=4). Scale bar = 50 µm. **(I)** Differentiation index (total nuclei in MF20^+^ myotubes to total nuclei) (***p<0.001, N=4). **(J)** Fusion index (percentage of total nuclei in myotubes with a range of number of nuclei to total nuclei) (*p<0.05, N=4). One-way ANOVA followed by Turkey’s post-hoc test and unpaired Student’s *t*-test.

Next, we induced myogenic differentiation in *Esrra* KD and control siNT myoblasts and measured the expression of early myogenic marker Myogenin (MyoG), and fusion index of differentiating myocytes on days 1 and 3 after induction of differentiation. ERRα protein expression was induced during myogenic differentiation in the control cultures but not in the *Esrra* KD myoblast cultures **(Figure 1D)** which confirms the KD efficiency of si*Esrra* and indicates that ERRα could have a role during myogenic differentiation. Interestingly, MyoG expression increased simultaneously with ERRα, with their protein levels reaching peak expression on day 2 after induction of differentiation in the control group. *Esrra* KD significantly reduced MyoG expression compared to the control **(Figure 1D)**. Transcript levels of these genes also exhibited the same expression pattern as the protein **(Figure 1E)**. We further examined MyoG expression by immunofluorescence staining after 1 day of induction of differentiation **(Figure 1F)**. We found that MyoG^+^ cells were lower in KD compared to the control group **(Figure 1F-G)**. Next, cell fusion was measured by immunostaining the cultures for myosin heavy chain (MyHC), a marker of the late stage of myogenic differentiation **(Figure 1H)**. Differentiation index was calculated as a percentage of nuclei within MyHC^+^ cells to total nuclei. On day 3 after induction of differentiation, the differentiation index was significantly reduced in the *Esrra* KD cultures compared to control cultures **(Figure 1I)**. Consistent with these results, quantification of fusion index, which is defined as the percentage of nuclei within MyHC^+^ cells at different subgroups of myotube size (1, 2 to 5, 6 to 9, greater than 10 nuclei myotubes) to total nuclei, showed there was a significantly lower ratio of myotubes containing 6 to 9 nuclei in *Esrra* KD vs. control cultures **(Figure 1J)**. Myotubes with greater than ten nuclei were absent in KD vs. the control cultures. These results demonstrate that ERRα plays an important role in regulating both proliferation and differentiation of myoblasts.

### 3.2. *Esrra* knockdown reduces angiogenesis and mitochondrial gene transcriptome during C2C12 myogenesis

Noting that ERRα transcriptional program in myogenic cells has not been decoded, we performed RNA sequencing in both proliferating and differentiating C2C12 myoblasts with or without *Esrra* KD. Compared to control cells, *Esrra* KD differentially downregulated 367 genes in proliferating myoblasts and 293 genes in differentiated myotubes **(Figure 2A and 3A)**. Gene ontology analysis for biological processes revealed the top gene sets downregulated in *Esrra* KD proliferating myoblasts **(Supplemental Figure S1)**. A cluster of genes associated with muscle cell proliferation and migration, regulation of vascular endothelial cell proliferation, and regulation of vasculature development are significantly downregulated in *Esrra* KD myoblasts compared to the control cells **(Figure 2B)**. In contrast, *Esrra* KD predominantly caused a reduction in gene sets related to mitochondrial function, ATP synthesis, and oxidative phosphorylation in differentiated myotubes **(Supplemental Figure S2 and Figure 3B)**.

**Figure 2.**
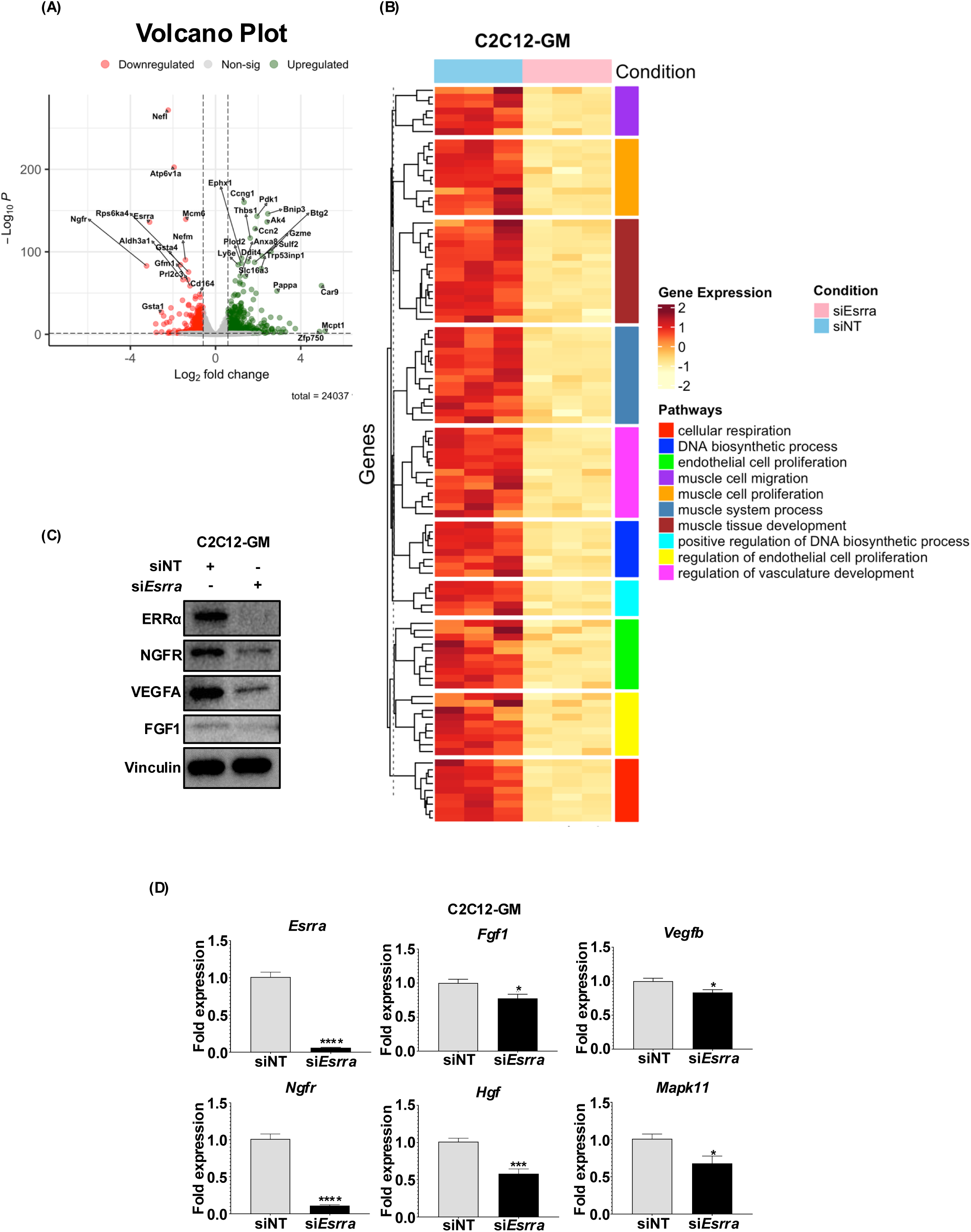
ERRα encoded transcriptional program in proliferating myoblasts. Following measurements were made in C2C12 cells treated with control scramble (siNT) or *Esrra* siRNA (si*Esrra*). **(A)** Volcano plot representing differentially expressed genes (953 genes) including 367 downregulated genes and 586 up-regulated genes in *Esrra* KD myoblasts compared to control with adjusted p<0.05 and |FC|>=1.5. (N=3). **(B)** Heatmap of major pathways downregulated in proliferating cells by si*Esrra* KD. **(C)** Representative western blot (WB) showing ERRα and angiogenic factors expression (N=3). **(D)** Quantification of Esrra and angiogenic gene expression (*p<0.05, ***p<0.001, ****p< 0.0001, N=6). Unpaired Student’s *t*-test.

**Figure 3.**
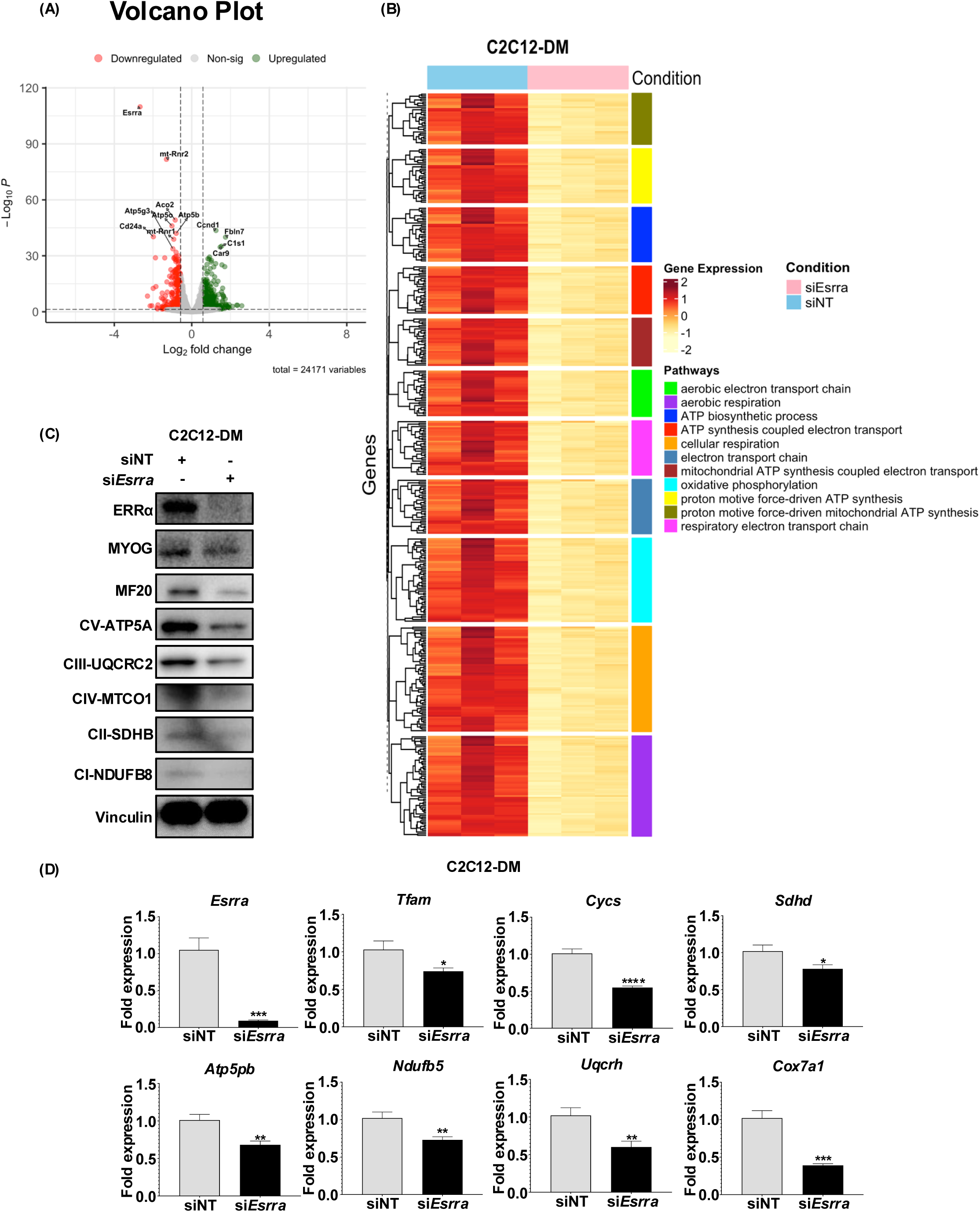
ERRα controlled transcriptional program in differentiated myoblasts. Following measurements were made in C2C12 cells treated with control scramble (siNT) or *Esrra* siRNA (si*Esrra*). **(A)** Volcano plot representing differentially expressed genes (775 genes) including 293 down-regulated genes and 482 up-regulated genes in *Esrra* KD myotubes compared to control with adjusted p-value < 0.05 and |FC|>=1.5. N=3. **(B)** Heatmap of major pathways downregulated in differentiating cells by si*Esrra* KD. **(C)** Representative western blot (WB) showing ERRα, myogenic marker and mitochondrial protein expression in *Esrra* siRNA-treated differentiated myotubes compared to control (N=3). **(D)** Quantification of *Esrra,* myogenic and mitochondrial gene expression in *Esrra* knockdown myotubes (*p<0.05, **p<0.01, ***p<0.001, ****p<0.0001, N=6). Unpaired Student’s *t*-test.

To validate RNA sequencing data, we measured gene and protein expression of angiogenic and mitochondrial targets, as well as other key myogenic genes in proliferating and/or differentiating C2C12 cells. Under proliferating conditions, we found that the protein expression of angiogenic growth factors (VEGFA and FGF1) was decreased in *Esrra* KD myoblasts **(Figure 2C)**. Likewise, a significant decrease in expression of angiogenic (*Vegfb*, *Fgf1*) and proliferation-related genes (*Hgf*, *Mapk11*) was observed in *Esrra* KD myoblasts compared to control cells **(Figure 2D)**. In differentiating cells, expression of OXPHOS proteins was decreased in *Esrra* KD differentiated myotubes compared to the control group **(Figure 3C)**. Furthermore, myogenic differentiation markers (MyoG and MyHC) were decreased in the KD cells. Consistent with protein levels, ERRα knockdown significantly reduced the mRNA levels of mitochondrial-related genes (e.g., *Tfam*, *Cycs*, *Sdhd*, *Uqcrh*, *Atp5pb*, and *Ndufb5*) **(Figure 3D)**. These results suggest that ERRα promotes myoblast proliferation and differentiation via transcriptional regulation of angiogenic, proliferative, mitochondrial and myogenic gene program.

### 3.3. ERRα promotes muscle regeneration in acute injury model

We next investigated the role of ERRα in muscle regeneration in muscle-specific ERRα overexpressing transgenic (TG) mice [31] and their age and sex-matched wild type littermates (WT). Acute muscle damage was induced by injecting BaCl_2_ into tibialis anterior (TA) muscles of TG and WT mice. Muscle regeneration was analyzed 4-, 7- and 15-days post-injury (dpi) **(Figure 4A)**. The non-injured TA muscle cryosections displayed normal myofibers with nuclei at the periphery of each myofiber, as shown by H&E staining **(Figure 4B)**. At 4 dpi, myofiber damage was universal, displaying many mononuclear cells and small regenerating myofibers within the injured area. Nascent myofibers were positive for embryonic myosin heavy chain (eMyHC), an early marker exclusively expressed in new regenerating myofibers **(Figure 4B, C)**. We found that there were significantly more eMyHC-positive myofibers in the regenerating TA muscle of TG compared to WT mice **(Figure 4D)**. Subsequently, the number of mononuclear cells decreased, and regenerating myofibers increased in size with lower expression of eMyHC at 7 dpi and 15 dpi. The size of myofibers in non-injured TA muscle was comparable between TG and WT mice **(Figure 4B)**. After injury, the regenerating myofibers were significantly larger in TA muscle of TG compared to WT mice at 7 dpi and 15 dpi **(Figure 4C, E)**, with myofiber size recovering to level of non-injured muscle in TG mice but not in WT by 15 dpi **(Figure 4E)**.

**Figure 4.**
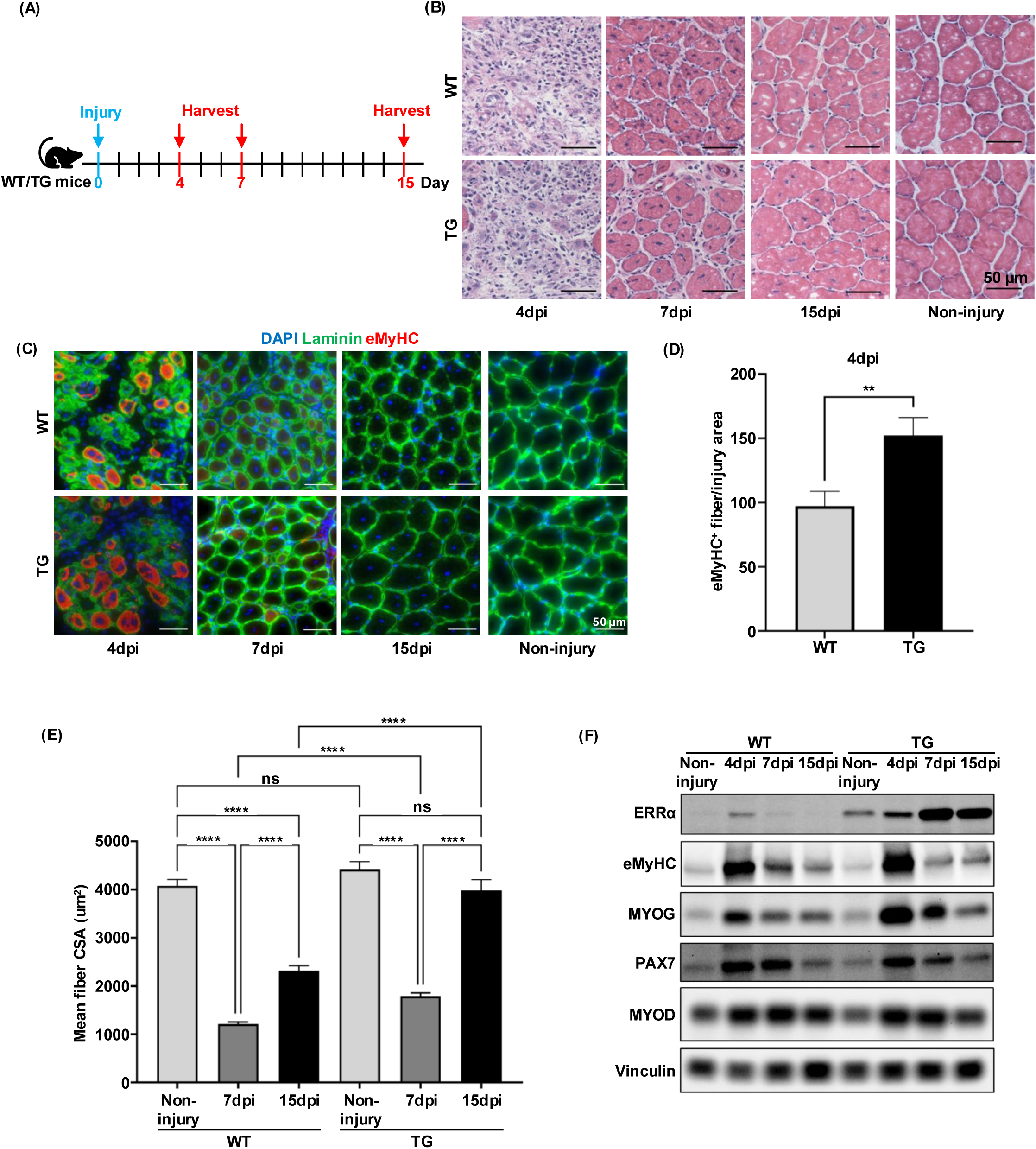
ERRα promotes muscle regeneration in BaCl_2_ injury model. **(A)** Experimental scheme for BaCl_2_-induced muscle injury and regeneration. Intramuscular injection of BaCl_2_ was carried out at day 0. TA were collected at 4, 7, 15 days post-injury (dpi). **(B)** Histological analysis of muscle regeneration by hematoxylin and eosin stain of TA cross-sections at 4, 7, 15 dpi compared to non-injured muscle (N=5). Scale bar = 50 µm. **(C)** Immunofluorescent staining of newly formed regenerating myofibers (eMyHC-red) at 4 dpi and centrally nucleated myofibers (Laminin = green, DAPI = blue) in the TA cryosections at 4, 7, 15 dpi compared to non-injured muscle. Scale Bar = 50 µm. **(D)** Quantification of newly regenerating myofibers (identified as eMyHC+ myofibers) derived from TA cross-section at 4 dpi (**p<0.01, N=5). **(E)** Quantification of regenerating myofibers size (identified as myofiber cross-sectional area) in TA cryosections from non-injured (N=2) and injured TA at 7 dpi (N=5) and 15 dpi (N=5) (****p<0.0001). **(F)** Representative western blot (WB) showing ERRα, eMyHC and Myogenin expression at 4 dpi and during regeneration at 7 dpi and 15 dpi (N=3). One-way ANOVA followed by Turkey’s post-hoc test and unpaired Student’s *t*-test.

These findings were confirmed at the level of myogenic marker expression in regenerating muscle. In WT mice, ERRα was induced at 4 dpi coinciding with the increased levels of eMyHC and myogenic markers such as Pax7, MyoD, and MyoG **(Figure 4F)**. Expression of eMyHC and MyoG were significantly higher in the TA of TG compared to WT mice, along with ERRα expression. However, there was no substantial difference in the induction of Pax7 and MyoD regenerating TA muscle of TG and WT mice. Collectively, these results demonstrate that ERRα overexpression promotes muscle regeneration in response to acute muscle injury.

### 3.4. ERRα enhances muscle angiogenesis and oxidative capacity in regenerating skeletal muscle

Based on the in vitro gene profiling data, we measured whether ERRα overexpression promotes muscle angiogenesis and mitochondrial activity in regenerating muscle. TA muscle cryosections were immunostained for CD31 (an endothelial cell marker) to measure angiogenesis/capillary density in non-injured and 15 dpi TA **(Figure 5A)**. The number of CD31-positive cells per myofibers was higher in TA muscle of TG vs. WT mice in non-injured muscles **(Figure 5B)**. At 15 dpi, we found that there was increased CD31 staining in both WT and TG muscle, which demonstrates elevated muscle angiogenesis during muscle regeneration. Strikingly, TG TA had significantly more CD31-positive cells per regenerating myofiber compared to WT mice. Therefore, muscle ERRα enhances new blood vessel formation and angiogenesis in regenerating muscle, which might partly contribute to increased muscle regeneration in TG vs. WT muscle.

**Figure 5.**
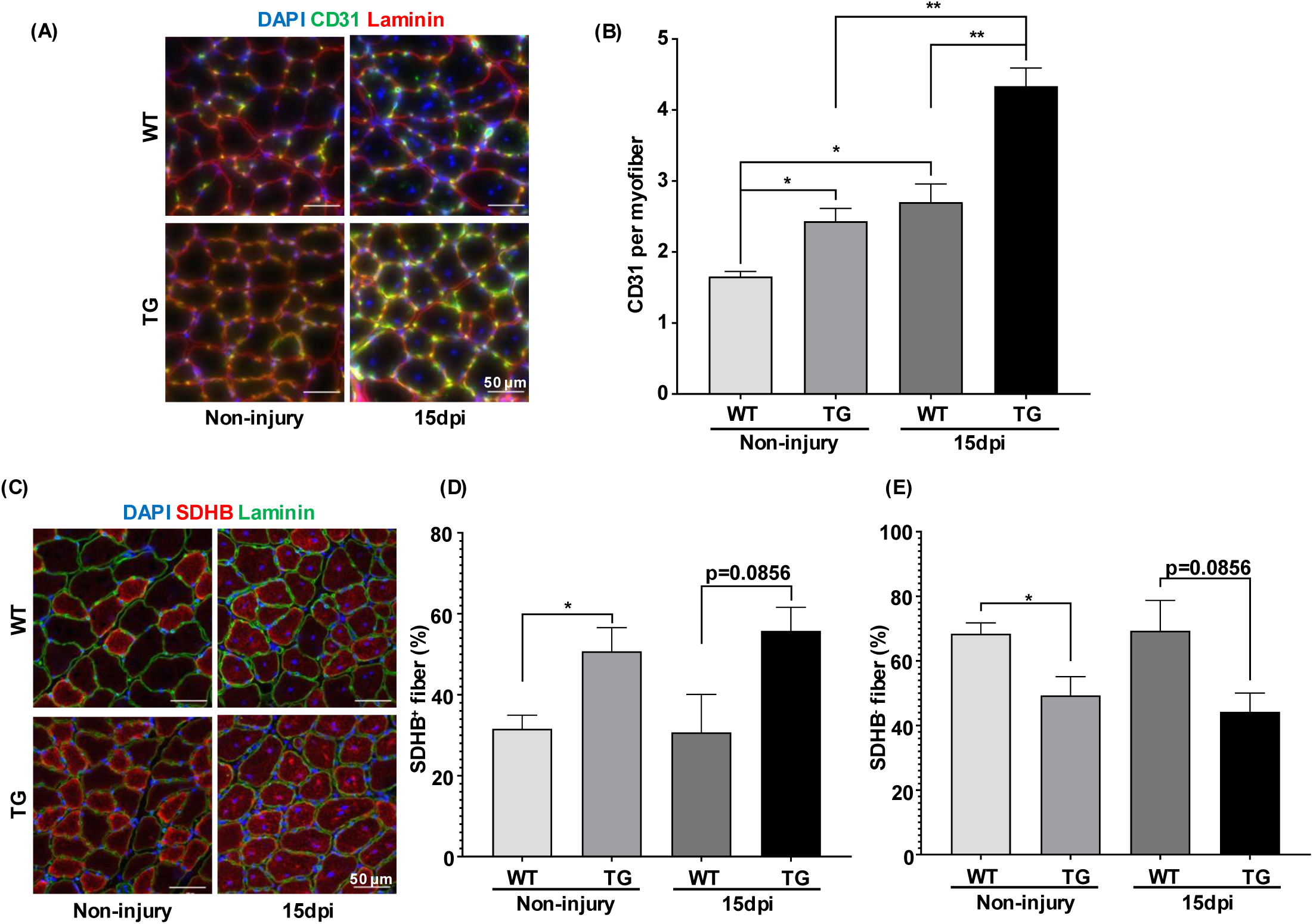
ERRα overexpression enhances angiogenesis and mitochondrial capacity in regenerating muscle. **(A)** Immunofluorescence characterization of capillaries (CD31^+^ endothelial cells) in non-injured and injured TA cross-sections at 15 dpi. Scale Bar = 50 µm. **(B)** Quantification of CD31^+^ endothelial cells per myofiber derived from non-injured (N=3) and injured TA cross-section at 15 dpi (*p<0.05, **p<0.01, N=5). **(C)** Immunofluorescence characterization of SDHB activity (SDHB^+^ = red) in non-injured and injured TA cross-sections at 15 dpi. Scale Bar = 50µm. **(D)** The percentage of SDHB^+^ myofibers from non-injury and injury TA cross-section at 15 dpi (*p<0.05, p=0.0856, N=3). **(D)** The percentage of SDHB^-^ myofibers from non-injured and injured TA cross-sections at 15 dpi (*p<0.05, p = 0.0856, N=3). One-way ANOVA followed by Turkey’s post-hoc test and unpaired Student’s *t*-test.

Next, we measured mitochondrial capacity in TA cross-sections. SDHB is an essential subunit of SDH, which is a mitochondrial enzyme complex involved in the tricarboxylic acid (TCA) cycle and the electron transport chain (ETC). SDHB staining is a reliable and well-known method to evaluate mitochondrial capacity. In non-injury muscles, there were significantly more SHDB-positive myofibers, and reciprocally fewer SDHB-negative myofibers in TA muscle of TG compared to WT mice **(Figure 5C-E)**. At 15 dpi, there was a trend towards increased SDHB-positive regenerating myofibers in TA of TG vs. WT mice, however without reaching statistical significance (p=0.0856). Consistent with these results, the number of SDHB-negative regenerating myofibers in TG mice injured TA muscle was lower than in WT injured TA. These results demonstrate that ERRα-mediated muscle regeneration is accompanied by angiogenic remodeling and mitochondrial biogenesis.

### 3.5. Impaired ERRα and target gene expression in dystrophic muscle stem cells (MuSCs)

MuSCs are crucial for regulating muscle repair after injury or disease-related damage and are typically exhausted in regenerative capacity in DMD. We first checked the status of ERRα signaling in dystrophic vs. non-dystrophic MuSCs. MuSCs were isolated from mdx and non-mdx mice using Magnetic-Activated Cell Sorting (MACS). MuSCs from mdx mice displayed a significant decrease in proliferation **(Figure 6A)**, with fewer EdU-positive cells compared to those from non-mdx mice **(Figures 6B–C)**. Notably, transcript levels of both *Esrra* and *Esrrg* are decreased in proliferating MuSCs from mdx mice. There was substantial downregulation of genes involved in angiogenesis and mitochondrial function in MuSCs from mdx mice, which correlated with the lower expression of *Esrra and Esrrg* **(Figure 6D)**. Myogenic differentiation capacity was also impaired in MuSCs from mdx mice, as measured by reduced staining for MyHC and a lower differentiation index **(Figures 6E–F)**. We also observed a decrease in *Esrra* and mitochondrial gene expression in myotubes derived from MuSCs of mdx mice upon induction of myogenic differentiation **(Figure 6G)**. Therefore, dysregulated expression of *Esrra* corelated with impaired proliferation and differentiation in MuSCs, suggesting that impaired ERRα signaling my contribute to impaired muscle regeneration, mitochondrial activity, and vascular supply in mdx mice.

**Figure 6.**
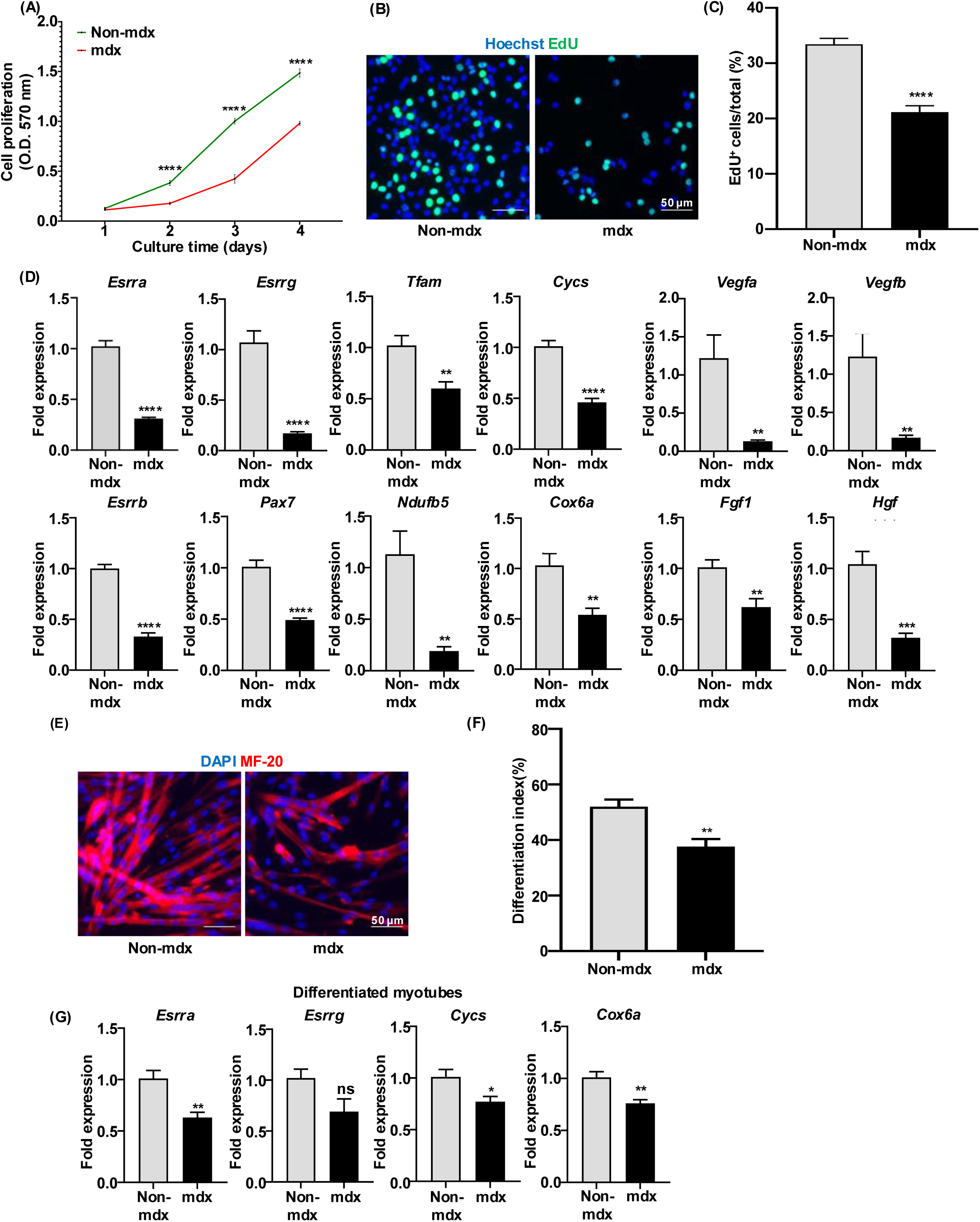
ERRα signaling in dystrophic muscle stem cells form mdx mice. Following measurements were performed in MuSCs isolated from the skeletal muscles of C57 control and mdx mice. **(A)** Proliferation in primary MuSCs (****p<0.0001, N=6). **(B)** Immunofluorescence characterization of S-phase cells (identified as EdU+ cells) in proliferating MuSCs (N=5). Scale Bar = 50 µm. **(C)** Percentage of S-phase cells (EdU+ cells) to total number of MuSCs (**p<0.01). **(D)** Quantification of *Esrra*, angiogenesis and mitochondria gene expression by qRT-PCR in proliferating MuSCs (**p<0.01, ***p<0.001, ****p<0.0001, N=6). **(E)** Representative images of immunostained myotubes (identified as MF20+ myotube) at 24 hours after myogenic differentiation of MuSCs (N=5). Scale Bar = 50 µm. **(F)** Quantification of differentiation index (identified as total nuclei in MF20+ myotubes to total nuclei) (**p<0.01, N=5). **(G)** Quantification of *Esrra* and mitochondrial gene expression by qRT-PCR in differentiated cells (*p<0.05, **p<0.01, N=6). Unpaired Student’s *t*-test.

### 3.6. ERRα overexpression ameliorates muscle damage and restores angiogenic and mitochondrial genes in mdx mice

To access the regenerative potential of ERRα in DMD, we generated mdx mice specifically overexpressing ERRα in the skeletal muscle (mdx-TG mice) by crossing the ERRα TG mice with the mdx mice **(Figure 7A)**. To test the effect of overexpressing ERRα in the mdx mice, we compared aged-matched inbreed C57Bl/6J non-mdx control mice, mdx mice, and mdx-TG mice. We first measured serum creatine kinase (CK) levels, a marker of myofiber damage, in these three groups. As expected, there is a significant rise in serum creatine kinase (CK) levels in mdx mice compared to non-mdx control mice, indicating more muscle damage **(Figure 7B).** ERRα overexpression in the skeletal muscles of mdx mice significantly reduced CK levels, indicating that overexpression of ERRα reduces muscle injury in mdx mice.

**Figure 7.**
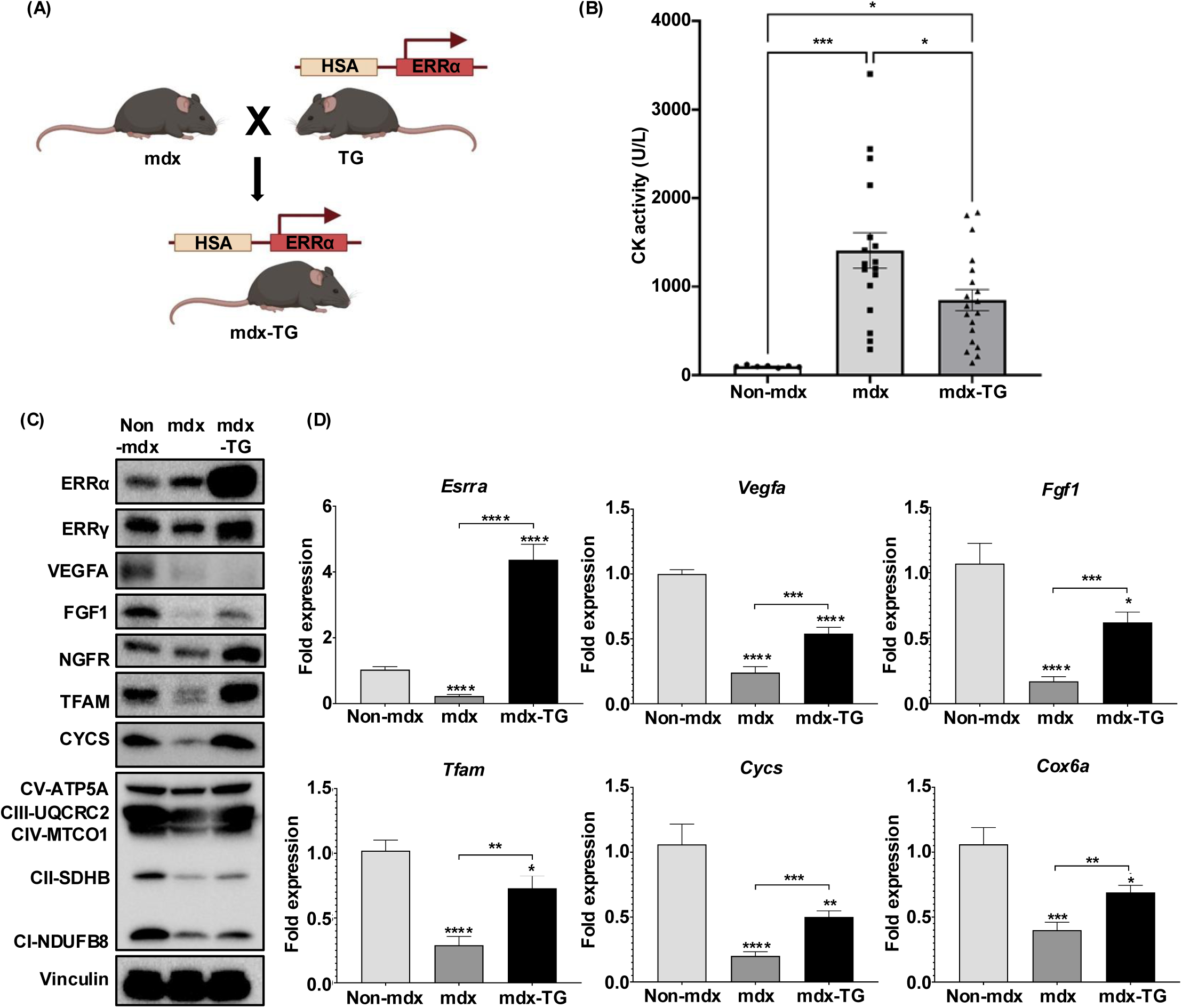
ERRα overexpression ameliorates muscle damage and restores angiogenic and mitochondrial gene expression in mdx mice. **(A)** Experimental breeding scheme to generate mdx and mdx mice overexpressing ERRα in the skeletal muscle (mdx-TG mice). Following measurements were made in C57Bl/6J controls, mdx and mdx-TG mice. **(B)** Quantification of serum creatine kinase (*p<0.05, ***p<0.001, N=7-19). **(C)** Representative western blot (WB) for ERRα, angiogenic factors and mitochondrial protein expression in TA muscle (N=3). **(D)** qRT-PCR analysis showing mRNA expression of *Esrra* and other genes linked to angiogenesis and mitochondria in TA muscle (*p<0.05, **p<0.01, ***p<0.001, ****p<0.0001, N=8). One-way ANOVA followed by Turkey’s post-hoc test and unpaired Student’s *t*-test.

Next, we compared the expression of ERRα/γ, as well as angiogenic and mitochondrial markers in TA muscle of non-mdx, mdx, and mdx-TG mice. Levels of ERRα and ERRγ protein were reduced in TA muscle of mdx compared to non-mdx mice **(Figure 7C)**. Further, protein levels of angiogenic factors (VEGFA, FGF1, NGFR) and mitochondrial factors (TFAM, CYCS, OXPHOS complexes) were reduced in the TA muscle of mdx mice compared to non-mdx mice. Interestingly, ERRα overexpression restored expression of ERRγ, as well as angiogenic and mitochondrial factors **(Figure 7C)**. Likewise, we found that the gene expression of *Esrra*, angiogenic genes (*Vegfa*, *Fgf1*), and mitochondrial genes (*Tfam*, *Cycs*, *Cox6a*) was suppressed in TA muscle of mdx mice, but was restored in mdx-TG mice upon ERRα overexpression **(Figure 7D)**. These results demonstrate that reduced levels of ERRs, mitochondrial, and angiogenic molecules in dystrophic muscles inversely co-relates with increase in serum CK levels. Whereas restoration of ERRs, metabolic, and angiogenic genes by ERRα transgene overexpression correlates with reduced serum CK levels equating to decreased muscle damage.

We next examined the effect of ERRα overexpression on muscle damage as a function of myofiber size in mdx TA in non-injured and BaCl_2_ injured muscle (15 dpi). The BaCl_2_ injury in mdx mice was used to determine whether acute injury applied in adult dystrophic muscle can be repaired by ERRα in real-time. We found that the baseline myofiber size was larger in mdx-TG compared to control mdx TA **(Figure 8A, B),** indicating the ERRα overexpression enhanced baseline muscle regeneration in mdx mice. After BaCl_2_ injury, we found reduced myofiber size in mdx TA muscle 15 dpi compared to non-BaCl_2_ injured mdx muscle. However, the size of regenerating myofibers in mdx-TG mice was significantly larger at 15 dpi compared to mdx control, demonstrating that ERRα can drive real-time muscle regeneration in dystrophic muscle after BaCl_2_ injury.

**Figure 8.**
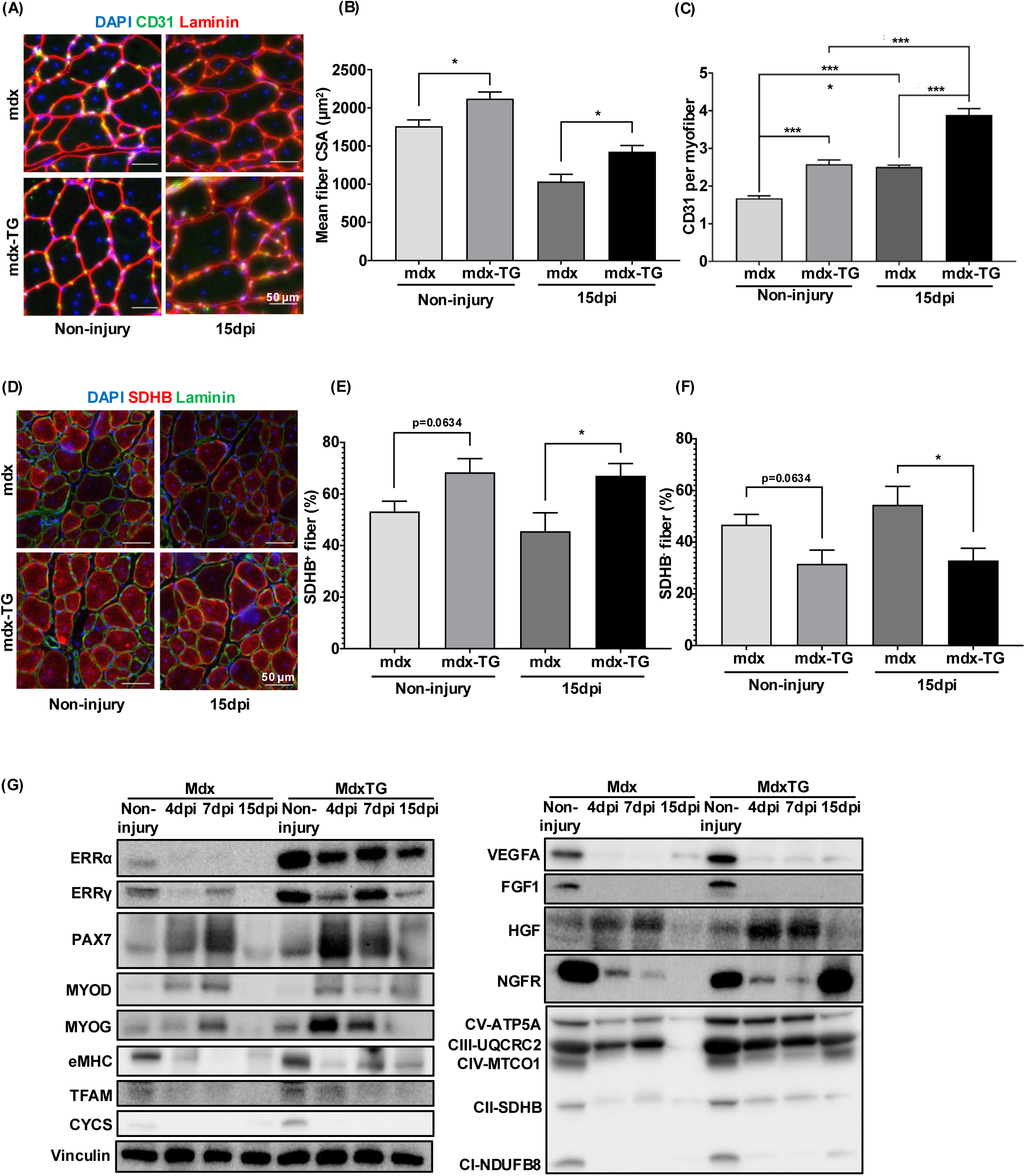
ERRα elevates myogenesis, angiogenesis and mitochondrial activity in mdx muscle. **(A)** Immunofluorescence staining of capillaries (identified as CD31^+^ endothelial cells) in non-injured and injured TA cross-sections at 15 dpi. Scale Bar = 50 µm. **(B)** Quantification of regenerating myofibers size (identified as myofiber cross-sectional area) derived from non-injury (N=3) and injury TA muscle cross-section at 15 dpi (N=5) (*p<0.05). **(C)** Quantification of CD31^+^ endothelial cells per myofiber derived from non-injury (N=4) and injury TA muscle cross-section at 15 dpi (N=4) (***p<0.001, ****p<0.0001). **(D)** Immunofluorescence characterization of SDHB activity (SDHB^+^-red) in non-injured and injury TA cross-sections at 15 dpi. Scale bar = 50 µm. **(E)** The percentage of SDHB^+^ myofibers from non-injured (N=4) and injury TA cross-sections at 15 dpi (N=5) (p=0.0634, *p<0.05). **(F)** The percentage of SDHB^-^ myofibers from non-injured (N=4) and injured TA cross-sections at 15 dpi (N=5) (p=0.0634, *p<0.05). **(G)** Representative western blot (WB) showing expression of myogenic and angiogenic factors and mitochondrial proteins in TA muscle (N=3). One-way ANOVA followed by Turkey’s post-hoc test and unpaired Student’s *t*-test.

We further supported the above experiments by measuring vascular and mitochondrial staining in mdx muscle. CD31 staining showed that angiogenesis/capillary density in mdx-TG TA was significantly higher than in TA muscle of mdx at both baseline and 15 dpi **(Figure 8A, C)**. While capillarity was significantly higher at 15 dpi in TA muscle of both mdx and mdx-TG mice compared to non-injured TA muscle, the mdx-TA showed maximal angiogenesis compared to all groups at 15 dpi. We also evaluated mitochondrial function by SDHB staining and found that there was an increase in SDHB-positive myofibers and a decrease in SDHB-negative myofibers in TA of mdx-TG mice compared to mdx mice in non-injury condition, without reaching statistical significance, nevertheless indicating a trend towards oxidative switch **(Figure 8D-F)**. However, overexpression of ERRα significantly increased the number of SDHB-positive myofibers while reciprocally decreasing the number of SDHB-negative myofibers in injured mdx-TG TA vs. mdx TA at 15 dpi **(Figure 8D-F)**. Therefore, ERRα ameliorates muscle damage and promotes muscle regeneration, accompanied by increased angiogenesis and mitochondrial biogenesis in the mdx mice.

To corroborate the effect of ERRα on muscle regeneration in mdx mice, we measured the protein levels of myogenic markers in non-injured and BaCl_2_-injured TA muscle on 4-15 dpi from mdx and mdx-TG mice. The protein levels of myogenic markers (PAX7, MYOG), early regenerating myofiber marker (eMHC), angiogenic and growth factors (VEGFA, FGF1, NGFR, HGF), mitochondrial proteins (TFAM, CYCS) were higher in TA of mdx-TG compared to mdx mice under non-injury conditions **(Figure 8G)**. ERRα overexpression overall enhanced the expression of myogenic, angiogenic, and mitochondrial proteins in TA muscle of mdx-TG vs. mdx mice at 4-15 dpi. These results agree with the effect of ERRα in C2C12 myoblast, which also show that ERRα activates myogenic, angiogenic, and mitochondrial gene program in the skeletal muscle facilitating the regenerative process.

## 4. DISCUSSION

We show that ERRα promotes myoblast proliferation and differentiation by activating angiogenic and mitochondrial biogenesis transcriptional programs, respectively, in a cell-autonomous fashion. Muscle-specific ERRα overexpression facilitates muscle regeneration after acute injury in mice, as well as mitigates dystrophic muscle pathology in mdx mice. ERRα and its downstream angiogenic and metabolic target genes are suppressed in proliferation and differentiation deficient dystrophic MuSCs, as well as in adult dystrophic muscles, likely contributing to worsening of dystrophy. We surmise that ERRα promotes muscle regeneration by stimulating angiogenesis and mitochondrial biogenesis in acute injury and murine model of muscular dystrophy.

ERRα controls adult muscle and exercise fitness by regulating fatty acid metabolism, mitochondrial biogenesis, angiogenesis, and vascularization [37; 38]. Here we demonstrate the muscle regenerative function of ERRα. We found that ERRα KD impairs myoblast proliferation and differentiation in C2C12 cells. Congruent effect of ERRα loss on myocellular differentiation was previously reported [39]. Our RNA-seq data in C2C12 cells goes further to provide new insights into transcriptional mechanisms underlying the regulation of myocellular proliferation and differentiation by ERRα. In proliferating cells, ERRα loss results in downregulation of genes linked to cell division/proliferation, and cell mobility/migration, but predominantly angiogenic growth factors. Several angiogenic growth factors (e.g. *Vegfa, Vegfb, Fgf1, and Ngrfr*) are downregulated in proliferating myoblast by ERRα KD. Angiogenic growth factors are linked to myoblast activation and proliferation. Vegfa is induced in activated satellite or muscle precursor cells, and it promotes myoblast activation/proliferation, inhibition of apoptosis, and survival of myotubes [40]. Vegfb stimulates myoblast proliferation and differentiation [9]. Hgf (hepatocyte growth factor) drives myoblast migration via activation of MAPK pathway [41]. Hgf also promotes satellite cell activation and regeneration [42; 43], primarily promoting muscle precursor proliferation over differentiation [44]. We postulate that ERRα-induced angiogenic growth factors promote myocellular proliferation via autocrine and/or paracrine effects.

In the differentiating cells, ERRα loss predominantly suppresses gene pathways linked with mitochondrial metabolism. Gene ontology analysis showed that genes associated with cellular respiration/oxidative phosphorylation/electron transport chain and ATP synthesis are targeted by ERRα. Differentiating myocytes increase their mitochondrial content to meet the energy demands of maturing myofibers [45–47]. Therefore, suppressed mitochondrial metabolism in differentiating cells due to ERRα loss is likely to impair myocellular differentiation. Other regulators of mitochondrial biogenesis/dynamics control myogenic differentiation. For example, PPR3 is a muscle-enriched protein that promotes myocellular differentiation via activation of the mitochondria [48]. Knockdown of mitochondrial dynamics regulator OPA-1 blocks C2C12 myoblast differentiation [49]. Myocellular aging or senescence induced by repeated passaging causes mitochondrial dysfunction, apoptosis, and impaired differentiation [50]. Using various complex-specific inhibitors of respiratory chain, Chabi et al, showed that pharmacological blockage of complex 1 and 2 impairs C2C12 myocellular differentiation into myotubes. The same study also showed that mitochondrial function could be critical for myoblast proliferation [47]. Further, master-regulators of mitochondrial biogenesis (e.g. PPARβ/δ, PGC1α, TFAM) promote myoblast differentiation [51; 52]. Therefore, ERRα could promote regeneration by driving proliferation or differentiation specific transcriptional effects on angiokine expression and mitochondrial biogenesis, respectively, in myogenic cells.

In BaCl_2_ injury model, muscle specific ERRα overexpression facilitates muscle regeneration. ERRα transgene expression is controlled by HSA promoter driving transgene expression in myoblasts and differentiating cells rather than MuSCs [53]; therefore, ERRα expression is unlikely to be impacted in MuSCs. Notably, ERRα overexpression simultaneously increased muscle angiogenesis and mitochondrial biogenesis in regenerating muscle, co-relating with the effects of ERRα we observed in cultured myogenic cells. Several angiogenic and mitochondrial regulators stimulate muscle regeneration in mouse injury models. Angiopoietin-1, which is an ERRα target gene in myocytes, stimulates muscle regeneration via muscle precursor cell activation and vascular regeneration via endothelial cell activation [54; 55]. Myogenesis by Angiopoietin-1 involves induction of MyoD [54; 55]. Adeno-associated virus mediated transfer of Vegfa to the skeletal muscle promotes muscle regeneration in models of chemical and ischemic injury, as well as in muscular dystrophy [56; 57]. Muscle-specific Vegfa knockout affects muscle architecture and likely affects muscle regeneration [58]. Similarly, Fgf1 promotes muscle precursor proliferation and muscle regeneration [59]. Therefore, ERRα could promote muscle regeneration by activating angiokine gene program in myogenic cells. It can be envisioned that ERRα drives transcriptional induction of angiokines in muscle precursors and/or mature myofibers around the injury area, which can drive myogenesis in autocrine or paracrine fashion in the transgenic mouse model. Likewise, mitochondrial biogenesis regulators (e.g. PGC1α, PPARδ), as well as direct mitochondria transfer have been shown to promote muscle regeneration in murine models [60–63]; therefore, it is likely that mitochondrial regulation is a part of ERRα-mediated muscle regeneration.

While the primary cause of myopathy in DMD is genetic mutation and loss of dystrophin, the pathology and severity are compounded by decreased MuSC function and regenerative capacity [5; 64–66]. Muscle specific ERRα overexpression in the skeletal muscle decreased the basal myopathy in mdx mice. The protective effect of ERRα arises from constitutive overexpression of the receptor since birth driving muscle regeneration. The real-time effect of ERRα on regeneration in dystrophic muscle was observed after reinjuring the mdx muscle with BaCl_2_, where ERRα boosted muscle regeneration after reinjury in mdx mice marked by post-injury increase in muscle fiber size recovery. Interestingly, we found that expression of angiogenic and mitochondrial metabolic genes is suppressed in dystrophic compared to the non-dystrophic muscle, a deficiency which is rescued by muscle ERRα overexpression in mdx mice. Therefore, ERRα-mediated mitigation of dystrophic muscle could partly involve recovery of angiogenic and metabolic gene program. Notably, ERRα expression and oxidative gene program are affected in various dystrophy models. ERRα and oxidative genes are repressed in the skeletal muscles of mdx mice and dystrophin/utrophin double knockout mice [25], which also exhibit structural defects in the mitochondria [67]. Expression of ERR sub-type ERRγ, which regulates mitochondrial biogenesis is downregulated in muscles of mdx mice [68]. ERRα and mitochondrial gene expression were repressed in muscle biopsies from DMD patients [69]. ERRα and mitochondrial downregulation were also observed in facioscapulohumeral muscular dystrophy [70]. Likewise, several studies have reported downregulation of angiokines, angiogenic dysfunction, and vascularization in dystrophic muscles of mdx mice [68; 71; 72].

Loss of dystrophin impairs stem cell polarity and asymmetric cell division decreasing the number of myogenic cells available for muscle regeneration [73]. Repeated injury in dystrophic muscle accelerates stem cell exhaustion and decline in regenerative capacity [74]. Compromised myogenic cell activation and regeneration are reported in mdx muscles in absence of repeated injury [75]. The myogenic defect is in part cell autonomous, as isolated dystrophic myoblast fail to proliferate and differentiate in culture [76]. MuSCs isolated from mdx mice have proliferation and differentiation deficits, and exhibit decreased ERRα/γ, angiogenic and mitochondrial biomarker expression compared to non-dystrophic muscle stem cells. Therefore, deficiency of ERRα in both dystrophic myofibers and in muscle stem cells might compound DMD pathology. Overall, ERRα activation promotes muscle regeneration in acute injury, as well as in mdx mice. The regenerative effect of ERRα is likely to involve replenishment of angiogenic and mitochondrial oxidative capacity in myogenic cells and/or adult muscle itself.

Estrogen-related receptors (ERRs) have emerged as master-regulators of metabolism and muscle physiological fitness [14; 17; 31]. Current study reveals multi-faceted role of ERRα in angiokine and metabolic regulation in regenerating muscle cells. While angiokines can act in autocrine and paracrine fashion, mitochondrial regulation could direct myogenic proliferation and/or differentiation. Our findings on ERRα and muscle regeneration raise exciting avenues for future investigations. ERRs could be expressed in MuSCs and regulate stem cell states (e.g. quiescence, activation, self-renewal). Noting the differential effects of ERRα in proliferative vs. differentiating muscle cells, it is likely that ERRs do so via recruiting distinct transcriptional complexes, chromatin modulation and/or post-translational receptor modifications under different myogenic states. Elucidation of these mechanisms could provide in-depth insights into transcriptional regulation of skeletal muscle regeneration. In DMD, investigating how ERR signaling is downregulated in dystrophic muscles could uncover new pathological mechanisms. Furthermore, in-depth transcriptomics of ERRα overexpression/activation in dystrophic muscle could provide insights beyond angio-metabolic regulation in mitigating muscle damage. A translational advantage of orphan nuclear receptors are the ligand binding pockets in the structure of these transcriptional factors, affording pharmacological modulation. Potent ERR pan-agonists have been recently developed [77], which have muscle boosting capacity. It will be interesting to measure the pharmacological effects of targeting ERRs in muscle regeneration. Similarly, the role of ERRs in muscle repair in dystrophies other than DMD should be explored. In summary, our work highlights the potential of targeting ERRs in muscle regeneration.

## Contribution

T.T.H.N. designed the experiments along with V.A.N, performed, analyzed and interpreted the data. Y.X.H., S.P., E.M., D.H.S performed experiments. C.C. and Z.Z. were responsible for RNA sequencing, data collection and analysis. A.K. was involved in data interpretation and editing of manuscript, and consolation with expertise in muscle degenerative diseases. V.A.N was responsible for hypothesis conception, planning of experiments, data analysis and interpretation and manuscript preparation with T.T.H.N.

