## Supplemental Information for "Estrogen-related receptor alpha promotes skeletal muscle regeneration and mitigates muscular dystrophy"

[1]Brown Foundation Institute of Molecular Medicine, McGovern Medical School, UTHealth, Houston, TX; [2]MD Anderson Cancer Center and UTHealth Graduate School of Biomedical Sciences, Houston, TX; [3]Department of Biology and Biochemistry, College of Natural Sciences and Mathematics, University of Houston, Houston TX; [4]Center for Precision Medicine, McWilliams School of Biomedical Informatics, UTHealth, Houston, TX; [5]Institute of Muscle Biology and Cachexia, University of Houston College of Pharmacy, Houston, TX; [6]Department of Pharmacological and Pharmaceutical Sciences, University of Houston College of Pharmacy, Houston, TX

[7]**Corresponding Author**

Vihang Narkar, Ph.D.
Professor
Center for Metabolic and Degenerative Diseases
Brown Foundation Institute of Molecular Medicine
McGovern Medical School
UTHealth
Houston, TX 77030


Keywords: skeletal muscle, dystrophy, regeneration, metabolism, angiogenesis

**Supplemental Figure S1.** Dot plot illustrating the top enriched biological processes of downregulated genes in ERRα KD proliferating myoblasts.

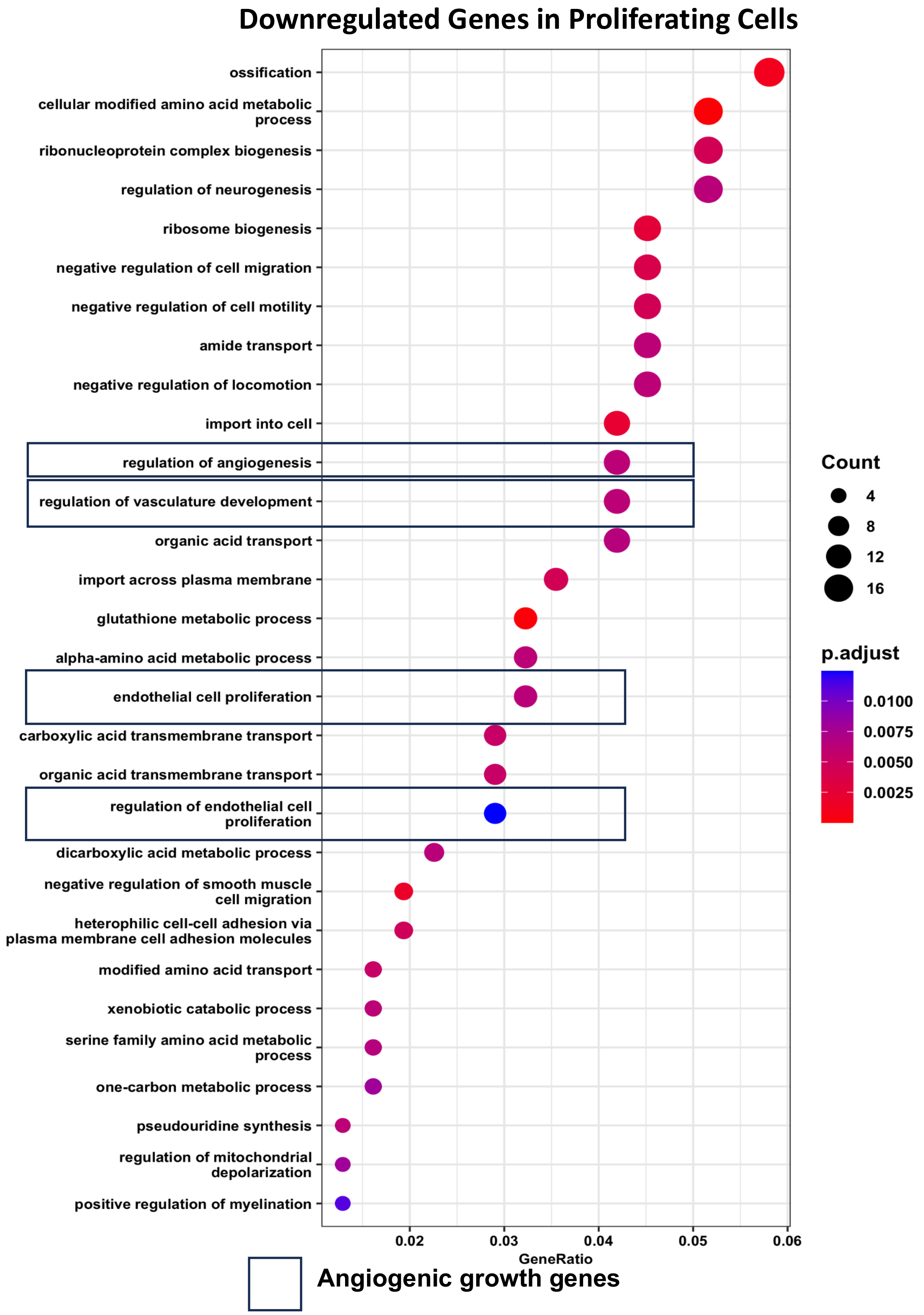

**Supplemental Figure S2.** Dot plot illustrating the top enriched biological processes of downregulated genes in ERRα KD differentiated myotubes.

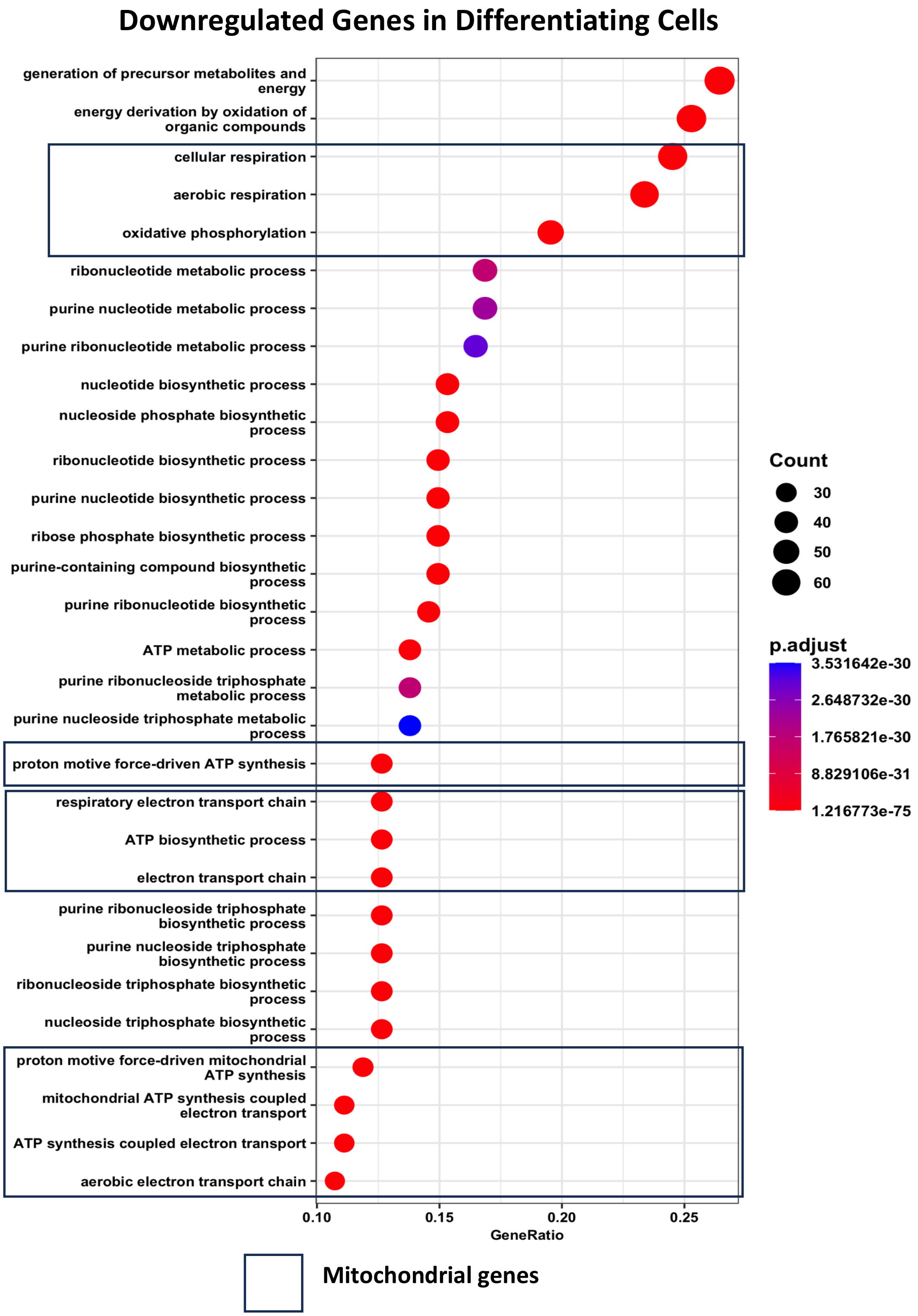

**Supplemental Table S1**. qRT-PCR mouse primers

| Gene | Primer | Sequence |
|---|---|---|
| *Esrra* | Fwd | CTCAGCTCTCTACCCAAACGC |
| | Rev | CCGCTTGGTGATCTCACACTC |
| *Cox6a* | Fwd | TCAACGTGTTCCTCAAGTCGC |
| | Rev | AGGGTATGGTTACCGTCTCCC |
| *Fgf1* | Fwd | GAAGCATGCGGAGAAGAACTG |
| | Rev | CGAGGACCGCGCTTACAG |
| *Ngfr* | Fwd | TGCCGATGCTCCTATGGCTA |
| | Rev | CTGGGCACTCTTCACACACTG |
| *Ppia* | Fwd | CCACCGTGTTCTTCGACAT |
| | Rev | CAGTGCTCAGAGCTCGAAA |
| *Hgf* | Fwd | CCTGACACCCCTTGGGAGTA |
| | Rev | TCCATAGGGACATCAGTCTCATTC |
| *Vegfb* | Fwd | TGCCATGGATAGACGTTTATGC |
| | Rev | TGCTCAGAGGCACCACCAC |
| *Mapk11* | Fwd | GCGGGATTCTACCGGCAAG |
| | Rev | GAGCAGACTGAGCCGTAGG |
| *Cycs* | Fwd | CCAAATCTCCACGGTCTGTTC |
| | Rev | ATCAGGGTATCCTCTCCCCAG |
| *Pdk4* | Fwd | AAGCAAAACACAAACACGAGTA |
| | Rev | CCCGGGTCATCCAACCA |
| *Sdhd* | Fwd | TGGTCAGACCCGCTTATGTG |
| | Rev | GGTCCAGTGGAGAGATGCAG |
| *Uqcrh* | Fwd | GTGGACCCCCTAACAACAGTG |
| | Rev | CGGGAAGACACGCGATTATCA |
| *Vegfa* | Fwd | GCACTGGACCCTGGCTTTAC |

| | | |
|---|---|---|
| | Rev | ATCGGACGGCAGTAGCTTCG |
| *Myog* | Fwd | CCTACAGACGCCCACAATC |
| | Rev | CCCAGCCTGACAGACAATC |
| *Tfam* | Fwd | ATTCCGAAGTGTTTTTCCAGCA |
| | Rev | TCTGAAAGTTTTGCATCTGGGT |
| *Atp5pb* | Fwd | AAAGTGCGTCTTGGGCTGATT |
| | Rev | CAAGCACATAAGGTCCTGTTACA |
| *Actb* | Fwd | ACGTTGACATCCGTAAAGAC |
| | Rev | GATCTTCATGGTGCTAGGAG |
| *Ndufb5* | Fwd | CAAGAGACTGTTTGTCGTCAAGC |
| | Rev | TGTTCACCAGTGTTATGCCAAT |
